# Macrophage-derived CCL24 promotes fibrosis and worsens cardiac dysfunction during heart failure

**DOI:** 10.1101/2024.07.12.603277

**Authors:** Preethy Parthiban, Fanta Barrow, Huy Nguyen, Fernando Neto, Kira Florczak, Haiguang Wang, Dogacan Yucel, Hong Liu, Micah Draxler, Erin Ciske, Gavin Fredrickson, Adam Herman, Marc E. Rothenberg, Samuel Dudley, Jop van Berlo, Xavier S. Revelo

## Abstract

Inflammation is a significant risk factor and contributes to cardiovascular disease by driving both adaptive and maladaptive processes. Macrophages are the most abundant immune cells in the heart and play an important role in the remodeling of cardiac tissue. We have previously shown an overall protective function of resident cardiac macrophages after pressure-overloaded injury. However, a subpopulation of resident macrophages also expresses high levels of the profibrotic CC motif chemokine ligand 24 (CCL24), suggesting a dichotomous role in pressure overload-induced cardiac remodeling. Here, we report that following transverse aortic constriction CCL24 knockout (CCL24 KO) mice have improved systolic function, cardiac wall enlargement, as well as increased myocyte surface area and hypertrophy, suggesting that CCL24 disrupts compensatory hypertrophy. TAC-operated CCL24 KO mice also displayed reduced fibrosis and diminished expression of fibrotic genes, implying a pro-fibrotic role for CCL24. Indeed, CCL24 induced the proliferation and activation of primary mouse fibroblasts in a process that required CCR3, the sole G protein-coupled receptor for CCL24. Correspondingly, selective ablation of CCR3 in fibroblasts improved cardiac function and ameliorated fibrosis following pressure overload. Administration of a CCL24 blocking antibody or a CCR3 antagonist both improved cardiac function in pressure-overloaded mice, highlighting the CCL24-CCR3 axis as a potential therapeutic target for heart failure. Finally, CCL24 deficiency improved cardiac function and ameliorated fibrosis during physiological aging. Overall, these results show that macrophage-derived CCL24 aggravates fibrosis via the CCR3 receptor, leading to impaired cardiac function in acute and chronic heart failure.

## INTRODUCTION

Cardiovascular disease (CVD) stands as the primary cause of mortality worldwide, affecting nearly half of the American population. In the US alone, the economic burden of CVD constitutes 12% of total health expenditures, surpassing costs linked to any other diagnostic group (American Heart Association). The relevance of inflammation in the progression of CVD has drawn considerable attention, prompting numerous clinical trials investigating anti-inflammatory therapies. However, these trials have predominantly failed, underscoring the imperative to comprehend the specific roles of immune cells within the heart^1, 2^.

Macrophages are the most abundant immune cells in the heart where they play pivotal roles in maintaining homeostasis and cardiac remodeling post-injury. As steady-state, cardiac resident macrophages (CRMs) perform tissue-specific functions such as efferocytosis, electrical conduction^3^, and mitochondrial homeostasis^4^. Adult cardiac macrophages can be distinguished into distinct subsets by their origin (monocyte- vs embryonic-derived) based on the expression of surface markers such as CCR2, TIMD4, LYVE1, and FOLR2^5, 6^. These subsets perform distinct functions in homeostasis and remodeling based on their ontogeny and/or the tissue microenvironment. Embryonically-derived CCR2^-^ CRMs are largely maintained through local proliferation unlike CCR2^+^ macrophages and were thought to mediate tissue-resolving functions^5^. The role of CRMs in disease, however, is poorly understood although recent work suggests that their function is dependent on the local pathophysiological niche. In a genetic mouse model of dilated cardiomyopathy, CRMs respond to mechanical stretch through the transient receptor potential vanilloid 4-dependent pathway to directly engage neighboring cardiomyocytes through focal adhesion complexes and promote adaptive hypertrophy^7^. In a mouse model of hypertension, CRMs secrete insulin-like growth factor-1 to regulate adaptive cardiac growth and maintain normal function^8^. In a mouse model of physiological aging and hypertension, CRM-derived interleukin (IL)-10 promotes fibrosis and diastolic dysfunction^9^. Overall, these findings show that macrophages are a heterogeneous population capable of performing specific roles in response to their microenvironment. However, the mechanisms by which CRMs regulate cardiac remodeling remain unclear.

CCL24 is a chemokine that activates cellular signaling by selectively binding to its sole receptor CCR3^10^. During inflammatory disease, CCL24 performs profibrotic and inflammatory roles in several tissues, including the kidneys, lungs, liver, and skin in animal models of disease. The neutralization of CCL24 using a monoclonal antibody decreases fibrosis and ameliorates disease in mouse models of primary sclerosing cholangitis^11^, metabolic dysfunction-associated fatty liver disease, experimental dermal and pulmonary fibrosis^12^. In the heart, macrophages are the major source of CCL24^13^, which is required for neonatal myocyte proliferation and cardiac regeneration shortly after birth^14^. However, the role of CCL24 in cardiac remodeling remains poorly understood. One study showed that the expression of cardiac CCL24 increases in patients with heart failure and cardiac fibrosis, while its blockade with a CCL24 antibody reversed hypertrophy and fibrosis in a mouse model of angiotensin II-induced heart failure^15^. Nevertheless, the underlying regulatory and effector mechanisms by which CCL24 regulates cardiac remodeling are unclear.

In this study, we investigated the role of CCL24 in the progression of acute pressure overload and chronic aging-induced heart failure. Utilizing CCL24-deficient mice, we demonstrate that CCL24 exacerbates cardiac dysfunction and remodeling following pressure overload. Using myocytes and primary mouse fibroblasts, we found a direct effect of CCL24 on impeding myocyte hypertrophy and activating fibroblasts through its receptor CCR3. Thus, we developed a mouse for conditional depletion of CCR3 and determined that specific genetic ablation of CCR3 in fibroblasts improves cardiac function and ameliorates fibrosis following pressure overload. Notably, pharmacological inhibition of both CCL24 and CCR3 rescues cardiac function following pressure overload injury. Finally, we determined that the pathological role of CCL24 in promoting fibrosis and cardiac dysfunction is preserved in chronic inflammation such as that underlying aging-induced heart failure. In conclusion, this study reveals that activation of the CCL24/CCR3 is an important immune mechanism that promotes cardiac pathology.

## RESULTS

### CRM-derived CCL24 promotes systolic dysfunction in response to pressure overload

We previously reported that CRMs are a heterogeneous population with key roles in the process of cardiac remodeling that follows pressure overload. Using single-cell RNA sequencing (scRNA-seq), we identified *Ccl24* as one of the cytokines specifically expressed by CRMs but not by their monocyte-derived counterparts^16^. As the source of CCL24 in the heart during cardiac remodeling has not been investigated, we re-clustered the macrophages in our published scRNA-seq dataset (GSE179276) and used known gene markers to calculate module scores for monocyte-derived macrophages (*Ccr2*, *Il1b*, and *H2-Ab1*) and CRMs (*Timd4*, *Folr2*, and *Lyve1*) (**Fig. 1A**). We then assessed the expression of *Ccl24* in sham and TAC-operated mice and confirmed that CRMs are the dominant cell type expressing *Ccl24* regardless of surgery (**Fig. 1B**). We also performed differential gene expression analysis between CCL24^+^ and CCL24^-^ macrophages and found a high expression of CRM markers in the CCL24^+^ cells while monocyte-derived macrophage markers were absent (**Fig. 1C**). Thus, our scRNA-seq data suggested that cardiac *Ccl24* expression is restricted to CRMs in normal and pressure overloaded conditions. To determine if the gene expression of *Ccl24* in the heart correlates with protein abundance, we measured CCL24 in cardiac lysates during early (1 week) and late (4 weeks) cardiac remodeling after TAC. We found that CCL24 protein levels increase twofold 1 week after TAC but return to baseline by 4 weeks after pressure overload (**Fig. 1D**), consistent with our previous findings showing a substantial increase in cardiac macrophages early after TAC and a return to baseline abundance at the 4-week timepoint^16^. Given that we previously identified IL-4 and IL-10 as upstream regulators of the CRM gene program^16^, we hypothesized that these factors synergically stimulate CCL24 production as observed in dermal resident macrophages^17^. First, we measured IL-4 and IL-10 in cardiac lysates and found that TAC increased their concentrations one week after the surgery (**Fig. 1E**). We next sorted total macrophages from uninjured mice, cultured them with IL- 4 or IL-10, and observed that the combined action of IL-4 and IL-10 stimulated a substantial production of CCL24 by cardiac macrophages while either cytokine alone failed to elicit CCL24 (**Fig. 1F**). This finding agrees with a previous report showing that IL-10 enhances the production of IL-4-induced CCL24 production by macrophages^18^. To confirm the requirement for IL-4 to produce CCL24 in vivo, we measured the amounts of cardiac CCL24 in IL4 KO mice and found that, compared with WT controls, IL4-deficient mice had decreased levels of cardiac CCL24 (**Fig. 1G**). Notably, CRMs express higher levels of IL-4 receptor than MoMFs (**Fig. 1H**), in agreement with the notion that resident macrophages are the primary cell type responsible for CCL24 production. Together, these results show that CRMs are the primary source of CCL24 in the heart during early cardiac remodeling in an IL-4-dependent manner.

**Figure 1.**
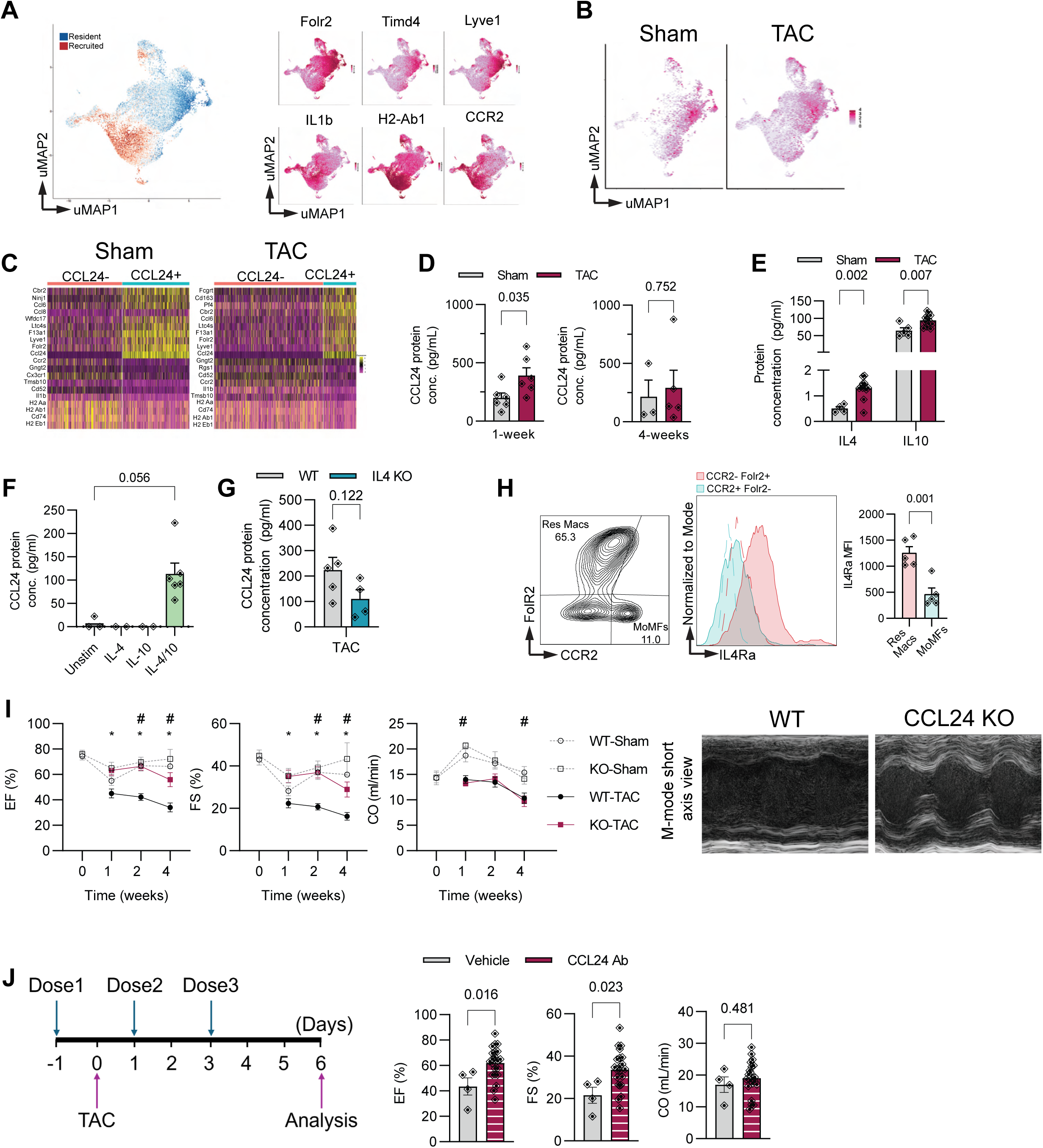
CCL24 upregulation following pressure overload. **A.** Uniform manifold approximation and projection (UMAP) projection of single cells clustered with identification of macrophage identity. Identification of cardiac resident macrophages (CRMs) and monocyte-derived macrophages (MoMFs) based on the expression of marker genes *Folr2* (Folate receptor beta), *Timd4* (T cell immunoglobulin and mucin domain containing 4), *Lyve1* (lymphatic vessel endothelial hyaluronan receptor 1), Il1β (Interleukin 1 beta), *H2-Ab1* (Histocompatibility 2, class II antigen A, beta 1), and *Ccr2* (C-C chemokine receptor type 2) are shown. **B.** Sham and transverse aortic constriction (TAC)–derived cells are plotted separately to visualize CCL24 abundance in both conditions. **C.** Heatmap of differentially expressed genes (DEGs) between CCL24+ and CCL24- macrophages. The identity of both subsets is indicated by top DEGs in sham and TAC conditions. **D.** CCL24 measured from whole cardiac lysates of sham- and TAC-operated hearts collected 1- or 4 weeks post-TAC. (n = 6, 6, 3, 5). **E.** IL-4 and IL-10 from sham- and TAC-operated hearts collected 1-week post-TAC (n = 3, 2, 2, 6). **F.** CCL24 secreted by isolated cardiac mϕs stimulated with 10ng/ml IL-4, IL-10 or both for 48 hours. (n = 3, 2, 2, 6). **G.** CCL24 measured from whole cardiac lysates of WT or IL4 knockout (IL4KO) TAC-operated mice collected 1-week post-surgery (n = 5,4). **H.** Flow cytometry analysis of IL-4 receptor on the surface of residential and recruited macrophages with gating strategy (left), mean fluorescence intensity (MFI) histograms (middle), and quantification (right) (n=5). **I.** Time course of systolic cardiac function at baseline, 1-, 2- and 4-weeks post TAC. EF – Ejection Fraction, FS – Fractional Shortening, CO – Cardiac Output. (Left – quantification; Right – representative M-mode echocardiography traces) (p value < 0.05 #- WT Sham vs WT TAC; *-WT TAC vs CCL24 KO TAC; + KO Sham vs KO TAC) (WT Sham: n = 11, 3, 5, 5; KO Sham: n = 11, 4, 6, 5; WT TAC: n = 8, 14, 11; KO TAC: n = 18, 15, 5). **J.** Schematic showing the timing of vehicle or CCL24 neutralizing antibody administration relative to transverse aortic constriction (TAC) surgery (left) and systolic cardiac function measured 1- week post-TAC following vehicle or CCL24 neutralizing antibody injections (right, n = 4, 21). Data pooled from 3 independent experiments.

To investigate the role of CCL24 on cardiac remodeling following pressure overload, we subjected CCL24 knock-out (CCL24 KO) and BALB/c wildtype (WT) control mice to either TAC or sham surgery. The depletion of CCL24 in the heart of CCL24 KO mice was confirmed by RT-PCR and ELISA (**Fig. S1A**). We assessed cardiac function in WT and CCL24 KO mice without any intervention or at 1, 2, and 4 weeks after sham or TAC surgeries by echocardiography. While there were no differences between WT and CCL24 KO mice without any intervention or after sham surgeries, CCL24 KO mice showed improved systolic function as evidenced by increased ejection fraction and fractional shortening at 1, 2, and 4 weeks after TAC, without any changes were observed in cardiac output (**Fig. 1I**). To determine the effect of pharmacological inhibition of CCL24 on cardiac function during pressure overload, we treated C57BL/6 mice with a CCL24 neutralizing antibody on days -1, 1, and 3 relative to TAC and performed echocardiography at the one-week timepoint. Using this approach, inhibition of CCL24 during the first week of cardiac remodeling led to a substantial improvement of systolic cardiac function to a level similar to that of CCL24 KO mice (**Fig. 1J**). To further confirm that CRMs are the source of pathological CCL24 after pressure overload, we transplanted bone marrow from WT and CCL24 KO mice into irradiated congenic recipient mice (**Fig. S1B**). As irradiation lethally depletes all immune cells including those in solid organs, the transplanted bone marrow cells exclusively replenished the cardiac MoMF compartment, while the resident macrophages remained depleted (**Fig. S1C**). After reconstitution, the chimeric mice were subjected to TAC, and cardiac function was assessed one week after pressure overload. There were no differences in cardiac function between recipients of WT or CCL24 KO bone marrow (**Fig. S1D**), suggesting that MoMFs from hematopoietic origin are not a source of functional CCL24 following pressure overload. Overall, these data show that genetic and antibody-mediated inhibition of CRM-derived CCL24 leads to a substantial improvement in cardiac function after pressure overload.

### CCL24 prevents compensatory hypertrophy following pressure overload

To determine if the improved cardiac function during CCL24 deficiency was associated with cardiac remodeling, we assessed cardiac chamber measurements and left ventricular hypertrophy in CCL24 KO mice. Compared with WT controls, CCL24 KO mice showed an increased left ventricular posterior wall (LVPWd) and decreased left ventricular internal diameter (LVIDd) 2 weeks post-TAC (**Fig. 2A**). Consistently, CCL24 KO mice had an increased heart-to-body weight ratio (HW/BW) one week after TAC (**Fig. 2B, left**), suggesting increased hypertrophy in the absence of CCL24. However, 4 weeks after TAC there was no difference in the HW/BW ratio (**Fig. 2B, right**), suggesting that hypertrophy was transient in CCL24 KO mice. There was no difference in the body weight between CCL24 KO and WT mice at either time point (**Fig. S2A**). In addition, the lung-to-body weight ratio was similar between WT and CCL24 KO mice post-TAC (**Fig. S2B**). To confirm hypertrophy at the cellular level, we assessed the cross-sectional area of myocytes by wheat germ agglutinin (WGA) staining one week after TAC and found increased myocyte area in CCL24 KO hearts (**Fig. 2C**). In contrast, myocyte size was similar between TAC-operated WT and CCL24 KO mice 4 weeks after surgeries, consistent with the lack of differences in HW/BW at this later timepoint (**Fig. S2C**). Gene expression analysis of cardiac tissue showed an increased expression of several hypertrophic genes such as *Nppa, Nppb, Serca2a,* and *Camk2d* in the heart of CCL24 KO mice one week after TAC (**Fig. 2D**). To determine if these effects were mediated by the direct action of CCL24 on myocytes, we treated fetal human ventricular cell line (RL-14) and neonatal rat ventricular myocytes (NRVMs) with 100ng/ml of recombinant CCL24. RL-14 cells showed a decrease in hypertrophic genes such *Nppa* and *Serca2a* following CCL24 stimulation (**Fig. 2E**). Additionally, both myocyte models showed a 15-20% reduction in cell size in response to CCL24 establishing its direct impact in impeding hypertrophic myocyte growth (**Fig. 2F**). Together, these results suggest that CCL24 worsens cardiac function and impedes compensatory hypertrophy during the adaptive phase of cardiac remodeling following pressure overload.

**Figure 2.**
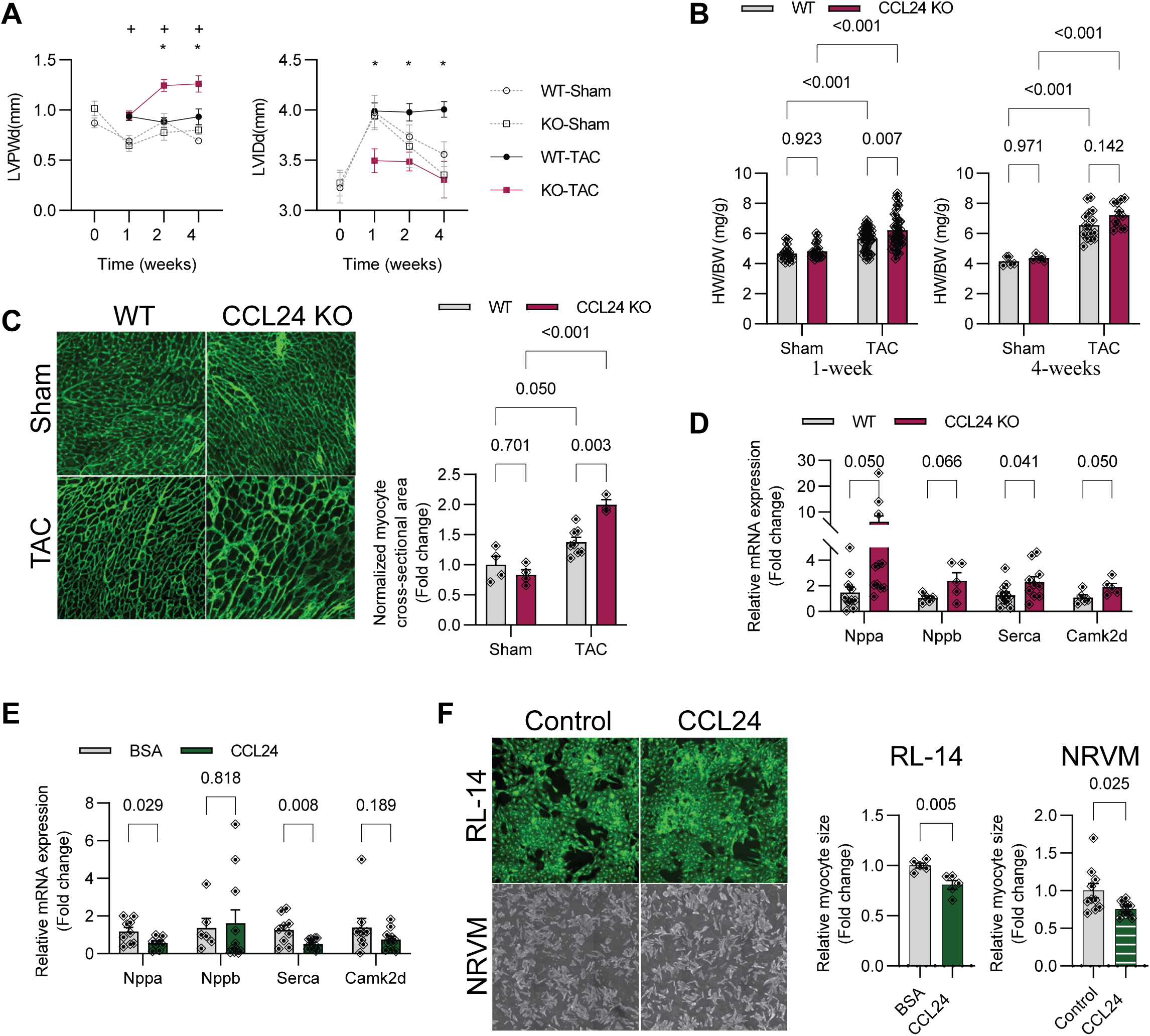
CCL24 worsens cardiac function following TAC injury. **A.** Time course of chamber measurements at baseline, 1-, 2- and 4-weeks post TAC. LVPWd – Left Ventricular Posterior Wall in diastole, LVIDd – Left Ventricular Internal Diameter in diastole (p value < 0.05 *-WT TAC vs CCL24 KO TAC; + KO Sham vs KO TAC) (WT Sham: n = 11, 3, 5, 5; KO Sham: n = 11, 4, 6, 5; WT TAC: n = 8, 14, 11; KO TAC: n = 18, 15, 5) **B.** Heart weight to body weight ratios (HW/BW) measured at 1- and 4-weeks post-TAC. (n = 21, 41, 21, 41). Data pooled from 3 independent experiments. **C.** Representative histological images (left) of WGA (wheat germ agglutinin; green) and quantification (right) of cross-sectional myocyte area from WT or CCL24 KO hearts 1-week post-TAC. **D.** Quantitative reverse transcription polymerase chain reaction (RT-qPCR) for atrial natriuretic factor (Nppa), brain natriuretic peptide (Nppb), sarcoplasmic/endoplasmic reticulum Ca2+- ATPase (Serca) and Calcium/Calmodulin Dependent Protein Kinase II Delta (Camk2d) on cDNA isolated from whole cardiac lysates from WT or CCL24 KO TAC-operated mice (**D**) or Human fetal ventricular cardiomyocyte, RL-14 cell line stimulated with 100ng/mL CCL24 for 48 hours (**E**). **F.** Representative images (left) of WGA staining of RL14 cell line (top) and bright field imaging of neonatal rat ventricular myocytes (NRVM) (bottom) stimulated with 100ng/mL of Bovine serum albumin (BSA) or CCL24 for 48 hours. Quantification of relative myocyte size normalized to a control group (right).

### CCL24 worsens fibrosis through activation of cardiac fibroblasts

To better understand the mechanisms by which CCL24 promotes cardiac dysfunction following pressure overload, we performed bulk RNAseq of cardiac tissues from WT and CCL24 KO mice one week after sham or TAC surgeries. Principal component analysis (PCA) showed that the most variation in the gene expression data was caused by TAC followed by CCL24 deficiency (**Fig. 3A**). A total of 333 and 357 genes were differentially expressed (FDR < 0.05) between WT and CCL24 KO mice in the sham and TAC conditions, respectively. To explore the biological relevance of these genes, we performed gene set enrichment analysis (GSEA). Although no baseline differences in cardiac function were detected in sham conditions (**Fig. 2**), CCL24 KO mice showed downregulation of the gene sets “Metabolism of Proteins” and upregulation of “Metabolism”, compared with their WT controls (**Fig. 3B, left**). While not statistically significant, CCL24 KO mice also showed downregulation of the “Immune System” gene set, likely as the result of the CCL24 deficiency. However, among TAC-operated groups, we found that “Collagen Formation”, “Extracellular Matrix Organization” and “Collagen Biosynthesis and Modifying Enzymes” gene sets were substantially downregulated in CCL24 KO mice (**Fig. 3B, right**). Notably, CCL24 KO cardiac tissue showed a profound downregulation of fibrotic genes such as *Col1a2*, *Adamts2*, *Pcolce*, and *Col1a1* which are part of the “Collagen Formation” and “ECM Organization” gene sets (**Fig. 3C**). We also mapped the DEGs between TAC-operated groups on “TGF-beta signaling” KEGG pathway, which predicted that most of its genes were suppressed in CCL24 KO tissues (**Fig. S3A**). We further compared the TAC-operated groups using pathway analysis and observed that the top enriched pathways were “Extracellular matrix organization”, “External encapsulating structure organization”, “Extracellular structure organization” and “Collagen fibril organization” were downregulated in CCL24 KO mice (**Fig. 3D)**. Given the strong anti-fibrotic signature during CCL24-deficiency predicted by GSEA and pathway analysis we mapped the key fibrotic genes in volcano plots and observed that *Tgfb2, Col5a1, Postn, Col14a1,* and *Mmp2* were substantially downregulated in the heart of TAC-operated CCL24 KO mice, compared with WT controls (**Fig. 3E**). However, a few ECM remodeling genes such as *Tgfb2* and *Timp1* were also downregulated in CCL24 KO under sham conditions (**Fig. 3E**). To confirm these results, we performed gene expression analysis by qPCR and found a substantial decrease in the expression of fibrotic genes such as *Col1, Acta2, Mmp2,* and *Tgfß* in cardiac tissue from CCL24 KO mice **(Fig. 3F)**. Thus, our unbiased gene expression analysis revealed that CCL24 aggravates cardiac fibrosis after pressure overload.

**Figure 3.**
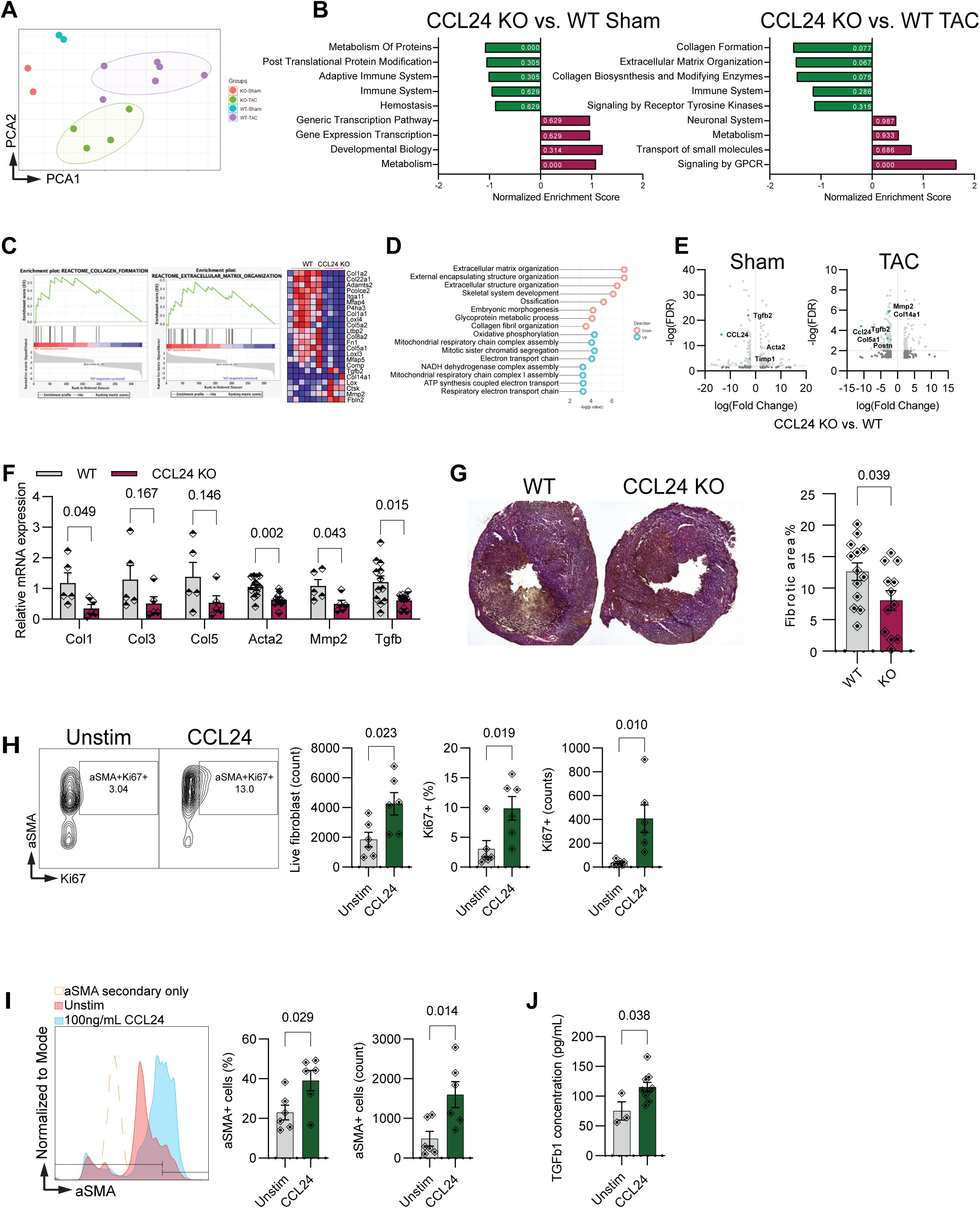
CCL24 worsens fibrosis through the activation of cardiac fibroblasts. **A.** 2-D Principal component analysis (PCA) of bulk RNA-sequencing from WT Sham (n = 2), CCL24 KO Sham (n = 2), WT TAC (n = 6), and CCL24 KO TAC (n = 4) groups 1-week post sham or TAC surgeries. **(B-C).** Top upregulated and downregulated gene sets detected by gene set enrichment analysis (GSEA) performed on differentially expressed genes between WT and CCL24 KO groups post sham (n = 333 genes) or TAC (n = 357 genes) surgeries (**B**). Enrichment plot for differentially regulated fibrotic pathways (**C, left**) and genes associated with the enrichment (**C, right**). **D.** Reactome pathway analysis for top upregulated and downregulated pathways between WT and CCL24 KO mice 1-week post-TAC. **E.** Volcano plot highlighting curated list of fibrotic genes differentially regulated in WT vs CCL24 KO mice in sham (left) and TAC (right) conditions. **F.** Quantitative reverse transcription polymerase chain reaction (RT-qPCR) for type I collagen (Col1), type III collagen (Col3), type V collagen (Col5), smooth muscle alpha (α)-2 actin (Acta2), matrix metallopeptidase 2 (Mmp2), and Transforming growth factor β (Tgfβ) on cDNA isolated from whole cardiac lysates from WT or CCL24 KO TAC-operated mice. **G.** Representative images (left) of Movat’s Pentachrome sections derived from WT or CCL24 KO mice post-TAC surgery. The bar graph (right) shows quantification of fibrosis (yellow staining, n=13, 13). Data pooled from 3 independent experiments. **(H-I).** Flow cytometry on primary mouse fibroblasts isolated from hearts and stimulated with 100ng/mL CCL24 for 24 hours. Quantification of Ki67+ fibroblasts (left) and representative plots of Ki67 vs. alpha-smooth muscle actin (aSMA) staining (**H, right**). Quantification of aSMA+ fibroblast (**I, left**) and mean fluorescence intensity (MFI) histogram of aSMA (**I, right**). **J.** ELISA for TGFb secreted by primary mouse cardiac fibroblasts stimulated with 100ng/mL of CCL24 for 48 hours.

Given the downregulation of fibrosis genes in CCL24 KO mice, we performed Movat’s Pentachorme and Picrosirius red staining of cardiac sections to determine their collagen content. Compared with WT controls, the hearts from CCL24 KO mice showed a 40% decrease in the fibrotic area at the one-week post-TAC timepoint (**Fig. 3G**). To determine if these effects were mediated by the direct action of CCL24 on cardiac fibroblasts, we treated primary mouse fibroblasts with 100 ng/ml of recombinant CCL24 and performed flow cytometry to assess fibroblast proliferation and activation. We found that the total number of fibroblasts, as well as the proportion and number of Ki67+ fibroblasts were increased in response to CCL24 stimulation (**Fig. 3H**). We also observed increased activation of fibroblasts as determined by a higher proportion and number of aSMA^+^ myofibroblasts in response to CCL24 (**Fig. 3I**). Since our KEGG pathway analysis predicted that the TGFB pathway was downregulated in CCL24 KO hearts, we tested whether CCL24 stimulates TGF*ß* production by cardiac fibroblasts. Compared with unstimulated controls, CCL24 increased the fibroblast-derived TGF*ß* in the supernatant (**Fig. 3J**), suggesting that CCL24 directly activates fibroblasts leading to enhanced TGF*ß* production.

### CCL24 promotes fibrosis through the CCR3 receptor

To better understand the cellular mechanism by which CCL24 promotes fibrosis, we next assessed the expression of CCR3, the sole receptor for CCL24, on cardiac cells under sham and TAC conditions. Flow cytometry analysis of the non-myocyte fraction showed a substantial expression of CCR3 on the cardiac fibroblasts (CFBs), followed by endothelial cells, and immune cells. In response to TAC, CCR3^+^ CFBs increased in number but not frequency while CCR3^+^ endothelial and immune cells remained unchanged (**Fig. 4A**). We also mined the expression of CCR3 in cardiac cells from a published dataset containing cellular gene expression analysis during pressure overload^19^ and found minimal CCR3 expression on cardiomyocytes and endothelial cells, but a robust increased in CCR3 gene expression of CFBs in response to TAC (**Fig. S4A**). To determine if CCR3 directly mediated the profibrotic actions of CCL24 in a TGF*ß*- dependent manner, we stimulated primary mouse fibroblasts with 100 ng/ml of CCL24 alone or in combination with blocking antibodies against CCR3 or TGF*ß*. Immunofluorescence assessment showed that CCL24 stimulated the expression of Periostin (Postn) as early as 24 hours after treatment followed by Collagen (Col1a1) and alpha-smooth muscle actin (αSMA) 48 hours after stimulation (**Fig. 4B**). Importantly, blocking CCR3 completely abrogated the stimulatory effect of CCL24 on Postn, Col1a1, and αSMA, indicating that fibroblast activation is mediated by the actions of CCL24 on CCR3 (**Fig. 4B**). The addition of a TGFß-blocking antibody also prevented the CCL24-mediated increase in Postn and αSMA, but not Col1a1 (**Fig. 4B**), suggesting that this process is at least partially mediated by the downstream actions of TGFß. To confirm these results using a secondary approach, we determined the expression of αSMA in cardiac primary fibroblasts by flow cytometry. We found that CCL24 treatment increased the expression of αSMA while blockade of CCR3 and TGFß reversed this effect (**Fig. S4B**). These findings provide substantial evidence that, following pressure overload, CCL24 directly activates cardiac fibroblasts via binding to its cognate receptor CCR3 and subsequent TGFß production.

**Figure 4.**
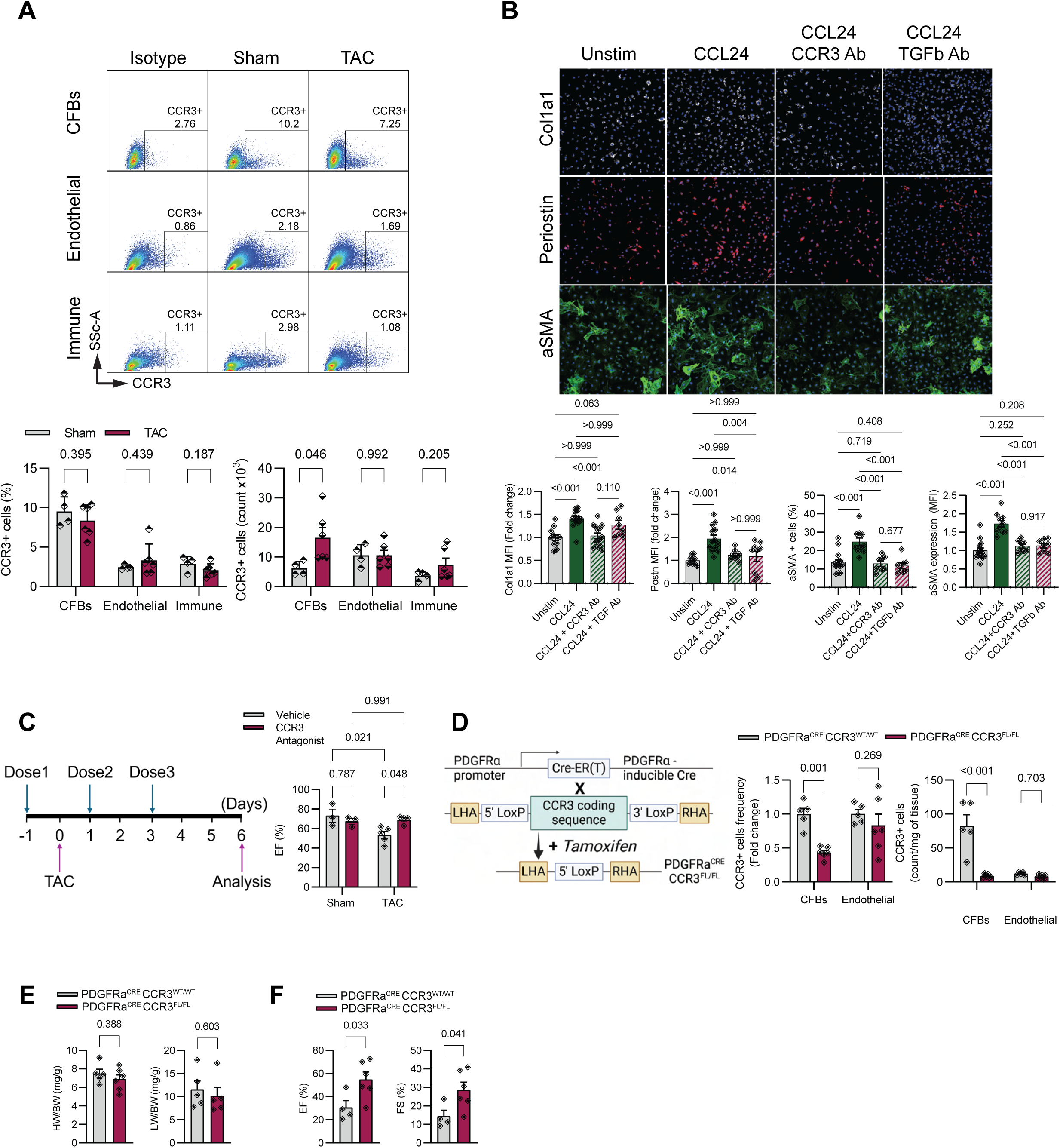
CCL24 promotes fibrosis through the engagement of the CCR3 receptor. **A.** Representative flow cytometry plots of cardiac non-myocytes showing the expression of CCR3 in the cardiac fibroblasts (CFBs), endothelial and immune cells from CCR3 isotype controls, sham or TAC-operated hearts (top). Quantification of CCR3+ proportions and cell counts in non- myocyte cell compartments (bottom, n = 4, 6). **B.** Representative images of fibroblasts stained for collagen-1 (white), periostin (red), αSMA (alpha-smooth muscle actin) (green), and DAPI (4’,6-diamidino-2-phenylindole) (blue) following treatment with 100 ng/mL of CCL24 alone or in combination with 0.5ug/mL of CCR3 or TGFb blocking antibody (n=4). Bar graphs show quantification of collagen-1, periostin and aSMA intensity, and aSMA+ cell percentage. **C.** Schematic showing the timing of vehicle or CCR3 antagonist administration relative to sham or transverse aortic constriction (TAC) surgery (left). Quantification of systolic function by ejection fraction (EF) in sham or TAC-operated mice receiving either vehicle or CCR3 antagonist (n = 3,3,5,4; right). **D.** Strategy for producing mutants with conditional deletion of CCR3 in fibroblasts. Mice homozygous for the CCR3 floxed allele were mated with transgenic mice expressing tamoxifen- inducible Cre recombinase under the control of a platelet-derived growth factor receptor alpha (PDGFRa) promoter, resulting in deletion of CCR3 in PDGFRa+ fibroblasts (left). The bar graph represents frequency and counts of CCR3+ fibroblasts and endothelial cells in PDGF^Cre^CCR3^wt/wt^ vs PDGF^Cre^CCR3^fl/fl^ mice 1-week post-TAC surgery. **E.** Heart weight to body weight (HW/BW) and lung weight to body weight (LW/BW) ratios in in PDGF^Cre^CCR3^wt/wt^ vs PDGF^Cre^CCR3^fl/fl^ mice 1-week post-TAC surgery (n = 5,5). **F.** Systolic cardiac function from M-mode short axis echocardiography of PDGF^Cre^CCR3^wt/wt^ vs PDGF^Cre^CCR3^fl/fl^ mice 1-week post-TAC surgery (n = 4, 6). EF – ejection fraction, FS – fractional shortening.

Next, we investigated whether the inhibition of CCR3 in vivo using the selective CCR3 receptor antagonist (SB328437)^20^ could improve cardiac function after pressure overload. Inhibition of CCR3 with the antagonist during the first week after TAC resulted in a 30% increase in ejection fraction (**Fig. 4C**), suggesting that systemic CCR3 blockade prevents the TAC-induced decline in systolic function. Given that CCL24 activates primary fibroblasts via CCR3, we wished to target CCR3 specifically in fibroblasts in vivo, but there were no transgenic mice available for this purpose. Thus, we used CRISPR/Cas9 to generate mice carrying a codon-optimized CCR3 coding sequence flanked by LoxP sites (**Fig. 4D, left**). Following backcrossing G0 animals to wild-type mice and validation via genotyping, we crossed the CCR3^fl/fl^ with the inducible PDGFRα^CreERT^ mice to deplete CCR3 specifically in cardiac fibroblasts^21^. Activation of Cre following 2 weeks of tamoxifen diet feeding led to the depletion of cardiac CCR3 expression specifically in fibroblasts while preserving expression in endothelial cells (**Fig. 4D, right**). Subsequently, we subjected tamoxifen-fed PDGFRα^CreERT^ CCR3^fl/fl^ and littermate PDGFRα^CreERT^ CCR3^WT^ controls to TAC and determined their cardiac function one week after surgeries. Despite no changes in the HW/BW and LW/BW ratios (**Fig. 4E**), the conditional deletion of CCR3 in fibroblasts led to a striking increase in cardiac function, as evidenced by increased ejection fraction and fractional shortening (**Fig. 4F**). These findings underscore the role of the CCR3 receptor on cardiac fibroblasts in promoting their activation and the development of fibrosis.

### CCL24-induced fibrotic remodeling is not associated with cardiac inflammation

As immune cell-driven inflammation is a key regulator of fibrotic remodeling in response to TAC, we next determined whether CCL24 worsens cardiac remodeling through alterations in cardiac and circulatory immune cells. First, we performed flow cytometry on circulatory and cardiac immune cells from WT and CCL24 KO mice at baseline without any intervention. Except for a mild decrease in Ly6C^low^ patrolling monocytes that survey the vasculature for pathogens in CCL24 KO mice, there were no substantial alterations in the frequency of circulatory eosinophils, neutrophils, and Ly6C^hi^ monocytes in CCL24-deficient mice (**Fig. S5A**). More importantly, the frequency of cardiac eosinophils, neutrophils, B cells, CCR2^+^ MoMFs, and CCR2^-^ CRMs was similar between WT and CCL24 KO mice (**Fig. S5B**), suggesting that CCL24 deficiency does not affect the local immune cell populations at steady-state. To determine if CCL24 deficiency alters the ability of immune cells to infiltrate the heart following pressure overload, we next used mass cytometry by time of flight (CyTOF) to quantify the major populations of immune cells in the heart of WT and CCL24 KO mice one week after TAC surgery. As we have previously reported^16^, macrophages were the most abundant immune cell population in the TAC-injured heart, followed by B cells, T cells, and neutrophils (**Fig. 5A-B**). However, CCL24 deficiency did not prevent the TAC-induced abundance of cardiac T cells, NK cells, B cells, neutrophils, eosinophils, and total macrophages (**Fig. 5A-B**), suggesting that CCL24 is not required for their accumulation following pressure overload. In separate experiments, we confirmed that CCL24 deficiency does not impact the accumulation of neutrophils, eosinophils, CCR2^+^ MoMFs, and CCR2^-^ CRMs by flow cytometry (**Fig. 5C**). Consistently, the expression of the inflammatory genes *Tnfa, Il1b* and *Il6* in whole cardiac tissue was similar between TAC-operated WT and CCL24 KO mice one week after surgery (**Fig. 5D**). Together, these data suggest that CCL24 does not promote cardiac dysfunction and fibrosis through decreased inflammation.

**Figure 5.**
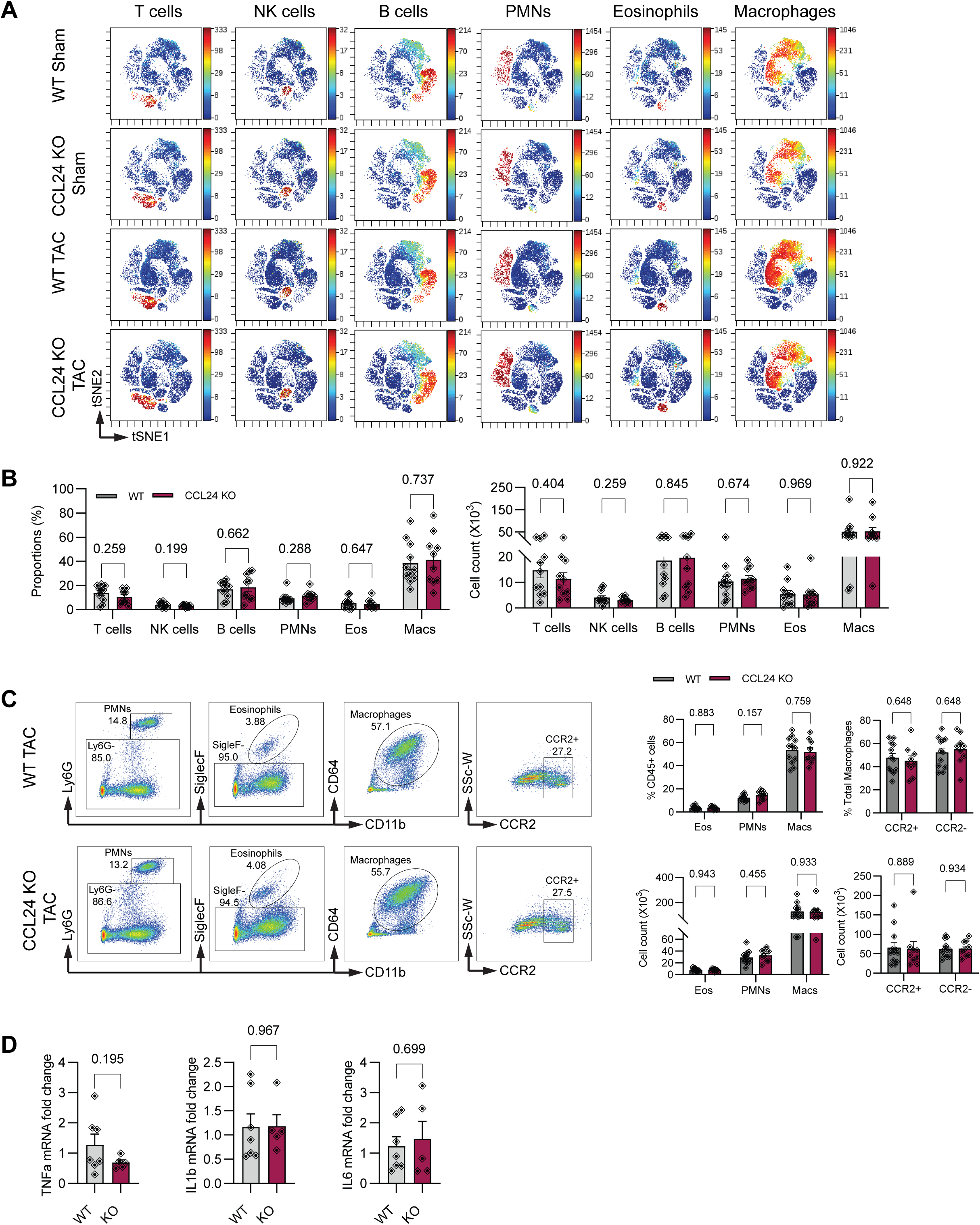
CCL24-induced fibrotic remodeling is not associated with cardiac inflammation. **A.** Cytometry by time-of-flight–based visualized stochastic neighbor embedding (ViSNE) plot showing the abundance of different immune cells and colored expression of CD3e (T cells), NKp46 (NK cells), B220 (B cells), Ly6G (PMNs), SiglecF (Eosinophils) and CD64 (Macrophages) in arbitrary units (AU) between WT-sham, WT-TAC, CCL24 KO-sham and CCL24 KO-TAC groups 1-week post-TAC. **B.** Quantification of immune cell frequency (left) and abundance (right) after TAC surgery in WT or CCL24 KO mice (n=11, 10). **C.** Representative flow cytometry plots (left) and quantification (right) of cardiac eosinophils (Eos), neutrophils (PMNs) and Macrophages (Macs) along with CCR2+ and CCR2- macrophages 1 week after TAC surgery in WT or CCL24 KO mice (n=12, 9). **D.** Quantitative reverse transcription polymerase chain reaction (RT-qPCR) for Tumor necrosis factor alpha (TNFa), Interleukin 1 beta (IL1b) and Interleukin 6 (IL6) on cDNA isolated from whole cardiac lysates from WT or CCL24 KO TAC-operated mice (n = 7, 5).

### CCL24 drives aging-associated cardiac dysfunction and fibrosis

Given our data showing a pathological role of CCL24 during the cardiac remodeling induced by an acute injury, we wished to determine if CCL24 promotes cardiac dysfunction and fibrosis in heart failure involving chronic inflammation. Interestingly, CCL24 has been identified as one of the canonical senescence-associated secretory phenotype SASP markers that predict aging-related senescence across tissues in humans^22, 23^. In the heart, CCL24 has been identified as a strong inflammatory marker of murine cardiac aging^24^. Thus, to determine if CCL24 has a direct role in aging-associated cardiac dysfunction, we first assessed the cardiac function of young (12- 14 weeks old) and old (85 weeks old) WT mice. Compared with young controls, old mice showed preserved systolic function but diastolic dysfunction indicated by increased E/A ratio with age (**Fig. 6A**). Furthermore, the internal diameter of the left ventricular chamber was increased in aged hearts suggesting enlarged chambers and dilation (**Fig. 6B**). As cardiac aging is associated with the development of cardiac fibrosis, we determined the expression of fibrotic genes and found that *Col1a1, Acta2* and *Tgfb* were significantly increased in aged hearts (**Fig. 6C**). Having established that old mice show the characteristic features of age-mediated cardiac dysfunction, we next measured the abundance of CCL24 in their cardiac tissue. Compared to young controls (12-14 weeks old), old WT mice (85 weeks old) showed a 20% increase in cardiac CCL24 levels (**Fig. 6D**). Notably, total cardiac macrophages isolated from old mice stimulated with IL-4/IL-10 had an increased capacity to produce CCL24, compared with young controls (**Fig. 6E**). To determine how aging alters the cardiac macrophage subsets, we performed flow cytometry to quantify CRMs and MoMFs. We found that the hearts from old mice had a shift in their macrophage subsets towards decreased CCR2^-^ CRMs while CCR2^+^ MoMFs were increased in proportion and number (**Fig. 6F**), suggesting that the increased CCL24 in the heart of old mice is the result of higher production of CCL24 per cell. Thus, these findings show that age-related cardiac dysfunction is associated with increased levels of CCL24 in the heart.

**Figure 6.**
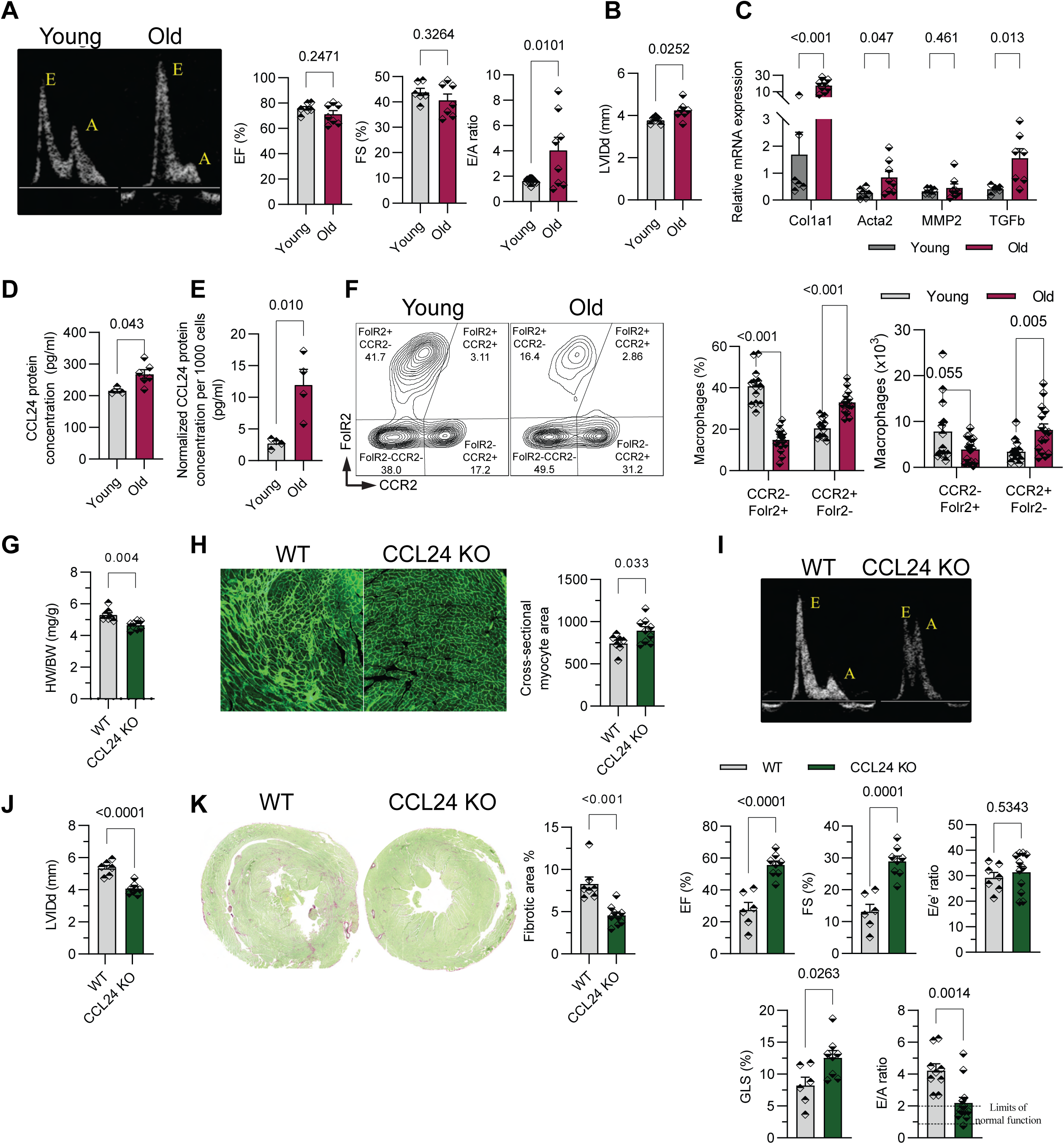
CCL24 drives aging-associated cardiac fibrosis and dysfunction. **A.** Representative E and A peaks from 4-chamber apical view echocardiography (left) and quantification of systolic function by ejection fraction (EF), fractional shortening (FS), and diastolic function measured by E/A peak ratio in wildtype young and old mice (right, n = 6, 7). **B.** Chamber dimension by left ventricular internal diameter in diastole (LVIDd) measured in wildtype young and old mice (right, n = 6, 7). **C.** Quantitative reverse transcription polymerase chain reaction (RT-qPCR) for type I collagen (Col1a1), smooth muscle alpha (α)-2 actin (Acta2), matrix metallopeptidase 2 (Mmp2), and Transforming growth factor β (Tgfβ) on cDNA isolated from whole cardiac lysates from young or old mice (n = 6, 7). **(D-E).** Enzyme-linked immunosorbent assay (ELISA) for CCL24 measured from whole cardiac lysates of wildtype young and old hearts (**D**, n = 3, 6) or isolated cardiac macrophages from wildtype young or old hearts stimulated with 10ng/mL IL-4/10 for 48 hours (**E**, n = 4, 4). **F.** Representative flow cytometry plots (left) and quantification (right) of residential (CCR2- Folr2+) and recruited (CCR2+ Folr2-) macrophages in wildtype young vs. old hearts (n = 13,15) Data pooled from two independent experiments. **G.** Heart weight to body weight ratios (HW/BW) measured in aged WT vs CCL24 KO hearts. (n =7,8) **H.** Representative histological images (left) of WGA (wheat germ agglutinin; green) and quantification (right) of cross-sectional myocyte area (n =7,8) **I.** Representative E and A peaks from 4-chamber apical view echocardiography (left) and quantification of systolic function by ejection fraction (EF), fractional shortening (FS), and diastolic function measured by E/e’, global longitudinal strain (GLS) and E/A peak ratio in aged WT and CCL24 KO mice. **J.** Chamber dimension by left ventricular internal diameter in diastole (LVIDd) measured in aged WT and CCL24 KO mice. **K.** Representative images (left) of Sirius red fast green (SRFG) sections derived from aged WT or CCL24 KO mice. The bar graph (right) shows the quantification of fibrosis (n=7, 8).

To determine if CCL24 has a causative role in aging-induced cardiac dysfunction and fibrosis, we aged WT and CCL24 KO mice for 60 weeks and assessed their cardiac phenotype. While the HW/BW ratio was lower in the CCL24 KO groups (**Fig. 6G**), the cross-sectional myocyte area was increased in CCL24-deficient mice (**Fig. 6H**), suggesting that CCL24-deficiency prevented the loss of cross-sectional volume in dilated myocytes. Compared with WT controls, aged CCL24 KO mice showed improved systolic and diastolic function with increased EF/FS, decreased E/A ratio, and increased global longitudinal strain (**Fig. 6I**). Furthermore, the internal LV diameter was reduced in CCL24 KO mice, suggesting decreased dilation and smaller chambers (**Fig. 6J**). Most notably, CCL24 deficiency during aging resulted in a nearly 2-fold decrease in cardiac fibrosis (**Fig. 6K**). Together, these data show that CCL24 promotes age-associated cardiac dysfunction and fibrosis.

## DISCUSSION

In the current study, we have demonstrated that CRM-derived CCL24 plays a pivotal role in promoting maladaptive remodeling by inhibiting compensatory hypertrophy and aggravating cardiac fibrosis. We have established a direct role for CCL24 on primary cardiac cells such as cardiomyocytes and fibroblasts. Mechanistically, we found that the CCL24 action of the CCR3 receptor on fibroblasts promotes their activation and proliferation through a TGFβ- independent pathways. While previous studies have identified a profibrotic role for CCL24 in several tissues^11, 12, 15^, here we show that the CCL24 signaling axis promotes cardiac dysfunction and fibrosis through the CCR3 receptor in cardiac fibroblasts. Thus, we provide evidence that CRMs not only play a crucial role in regulating pro-repair fibroblasts, but they can induce pro-fibrotic myofibroblasts primarily through the secretion of CCL24.

Here, we identified that elevated cardiac IL-4 and IL-10 are upstream triggers for CRM-derived CCL24 production. This finding is consistent with a previous study showing that dermal macrophages require the synergistic stimulation of IL-4 and -10 for CCL24 secretion during infection^17^. Similarly, the anti-inflammatory cytokine IL-10 secreted by CRMs has been shown to promote cardiac fibrosis and diastolic dysfunction^9^. Importantly, IL-4 is elevated in patients with heart failure and has been shown to promote cardiac fibrosis following pressure overload in pressure-overloaded mice^25^. In the context of angiotensin-induced hypertension, IL-4 promotes myocardial interstitial fibrosis and macrophage accumulation^26^. While the signals that determine the fibrogenic function of CRMs are unclear, excessive and chronic activation of type 2 responses that ameliorate inflammation can lead to the development of pathological fibrosis^27^. Type 2 immunity, characterized by increased production of the cytokines IL-4 and IL-13, signals macrophages to resolve inflammation and restore tissue homeostasis^28^. Although type 2 immunity exhibits many host-protective functions such as suppression of excessive inflammation and tissue regeneration, excessive and chronic activation of IL4 and IL13 can lead to pathological fibrosis.

Cardiac fibrosis is evolutionarily conserved given the protection and structural stability it provides during the early response to injury. In non-ischemic injuries such as pressure overload, there is an initial reactive interstitial fibrosis to adapt to the increased pressure gradient against which the cardiac muscle works, however, persistent fibrosis replaces necrotic cardiomyocytes and turns maladaptive, leading to stiffening of the heart^29^. Macrophages are well-known to contribute to fibrosis through the secretion of cytokines, growth factors, and their transdifferentiation into myofibroblasts^30, 31^. However, given their heterogeneity, different macrophage subpopulations can exert a bidirectional regulation of cardiac fibrosis^32^. We have shown that depletion of CRMs aggravates fibrosis and prevents angiogenesis leading to dysfunction in a TAC-induced pressure-overload model^16^. However, in rats fed with a high-salt diet or exposed to TAC injury, depletion of CRMs through clodronate liposomes led to reduced cardiac hypertrophy and fibrosis and led to improved function^33^. One possibility is that the fibrogenic role of CRMs is driven by the plasticity of their phenotype during cardiac repair. During the resolution of inflammation, CRMs are needed to remove apoptotic cells in a process that requires a phenotypic switch from a pro-inflammatory to an anti-inflammatory and/or reparative phenotype^34^. However, as CRMs aid in resolving inflammation via secretion of IL-4, IL-13, IL-10, and TGF-β1 that promote repair, unregulated production of these factors leads to pathological fibrosis^35^.

Interestingly, we found that CCL24 does not promote the recruitment of immune cells or alter the proportion of cardiac macrophage subsets during pressure overload, suggesting that CCL24 is not involved in immune cell trafficking or macrophage activation. Considering that CCL24 was originally discovered as an eotaxin that promotes the migration and activation of eosinophils^36^, we were surprised by this finding. One explanation is that the recruitment of eosinophils in CCL24 KO mice following pressure overload is compensated by the action of other eotaxins such as CCL11 that bind the CCR3 receptor^37^. Future work is needed to determine whether CCL11 has a differential or redundant role during cardiac remodeling. Another caveat of the current study is that the CCL24 KO mouse was developed on the BALB/c background, leading to potential inherent strain differences in cardiac remodeling compared with C57BL6 mice^38^. However, we found consistent cardioprotective and anti-fibrotic roles for the antibody-mediated CCL24 inhibition and genetic CCR3 depletion performed in C57BL/6 mice, suggesting that the different strains used in our study are not confounding factors.

Overall, here we show that CRM-derived CCL24 is a key cytokine that worsens fibrosis in pressure overload by stimulating cardiac fibroblasts through the receptor CCR3. Mechanistically, type 2 cytokines such as IL-4 and IL-10 promote the production of CCL24 by CRMs that instigate fibrosis following pressure overload. Importantly, we provide evidence that CCL24 directly activates cardiac fibroblasts via binding to its cognate receptor CCR3 in vivo. These data highlight an important mechanism by which macrophage-derived CCL24 promotes cardiac dysfunction and fibrosis during cardiac remodeling.

## MATERIALS AND METHODS

### Animal Procedures

All animal procedures were approved by the University of Minnesota Institutional Animal Care and Use Committee. C57Bl/6j (000664) and wildtype BALB/c (000651) animals were obtained from the Jackson laboratory. Whole-body CCL24 KO mice were gifted by Rothenberg lab at the Cincinnati Children’s Hospital. Cardiac pressure overload was induced via TAC as previously published. Briefly, 8-10-week-old animals were intubated and anesthetized with inhaled 2.5% isoflurane. A left parasternal incision was performed until the third rib. The lobes of the thymus were separated to expose the transverse aorta. A blunt 26/27G needle was tied to the transverse aorta between the brachiocephalic and left carotid arteries using a 6-0 silk suture (Ethicon K889H). The needle was removed, the thoracic wall was sutured with a 4-0 silk suture (Ethicon 683G), and the skin was closed with tissue glue (GLUture). Sham-operated animals underwent the same treatment, except for ligating the aorta. The surgeon was blinded to the genotype/treatment. Cardiac pressure overload was verified by Doppler imaging of the maximum velocity of blood flow across the constriction (VEVO 2100 and VEVO F2, Fujifilm). Cardiac dimension and function were analyzed using M mode imaging on both short and long axes and 4-chamber apical images. Mice were randomly assigned to sham or TAC surgery and male and female mice were used for all assays. To block CCL24 in vivo, we administered 3 doses (100 uL) of either vehicle control or 50 ug of CCL24 neutralizing antibody (MAB528, R&D systems) via intraperitoneal injection before TAC and during the first week of recovery. To inhibit CCR3, we administered 3 doses (100 uL) of either vehicle control or 50 ug of the CCR3 antagonist (SB328437, Tocris) via intraperitoneal injection before TAC and during the first week of recovery. Cages were randomly assigned to treatment groups.

### Bulk RNA Sequencing

Total RNA was extracted from cardiac tissue using the RNeasy Plus Mini kit (Qiagen). Illumina sequencing libraries were created using the SMARTer Stranded RNA Pico Mammalian V2 kit (Takara Bio) and sequenced on a single lane in a NextSeq 550 instrument (Illumina) using the 75bp paired-end Mid Output Mode. Differential gene expression analysis including principal component analysis (PCA) and pathway analyses was performed using edgeR (Bioconductor). Gene set enrichment analysis was conducted using GSEA software^39^.

### ELISA

Macrophages were labeled using anti-F4/80 microbeads and purified using LS columns and a MACS separator (Miltenyi Biotech). Purified macrophages from one heart were seeded per 96- well plate and were stimulated with 10 ng/mL of IL-4 and IL-10 for 48 hours. The supernatant was collected for downstream ELISA. For protein extraction from whole tissue, 20mg of heart pieces were weighed into an extraction buffer containing 1 mL of RIPA buffer and 1x cOmplete, EDTA-free Protease Inhibitor Cocktail (04693132001, Millipore Sigma). Tissues were homogenized using a bead rupture homogenizer. Samples were diluted 5X in assay buffer and assayed for ELISA. CCL24 was quantified using the R&D Biosystems capture and detection antibody pair (MAB528, BAF528). IL-4 and IL-10 were analyzed using the ELISA Max Deluxe set (431104) and Standard set (431411) respectively.

### Isolation of cardiac immune cells

Cardiac non-cardiomyocytes were isolated from the ventricular myocardium according to our previously published protocol^16^. Hearts were flushed with ice-cold PBS to remove red blood cells, minced into a slurry, and digested in Hanks Balanced Salt Solution (Corning 21022CM) with 2.4 U/ml Dispase II (Roche, 04942078001), 0.1% collagenase B (Roche, 11088831001), 10mM Hepes, and 2.5mM CaCl2for 30 min in a 37°C water bath. The suspension was triturated and strained through a 70μm cell strainer, centrifuged followed by red blood cell lysis (BioLegend, 420302), and strained through a 40μm cell strainer. Isolated non-cardiomyocytes were then analyzed by mass cytometry and flow cytometry using immune cell-focused panels.

### Mass cytometry

Isolated cardiac non-cardiomyocytes were stained with 5μM cisplatin (Fluidigm 201064) to discriminate between viable and dead cells. Cisplatin staining was quenched with Maxpar cell staining buffer (Fluidigm 201068). Non-specific binding was blocked with TruStain FcX Plus (Biolegend 101320). Staining for cell surface markers was performed with 0.5μg metal-conjugated primary antibodies (Major Resources Table) for 30 min at 4°C. Antibodies that were not available pre-conjugated were purchased and conjugated using a Maxpar labeling kit (Fluidigm 201300). Cells were fixed with 1.6% methanol-free Formaldehyde (Thermofisher) and incubated with 0.5μM intercalator solution in fix and perm buffer (Fluidigm 201192A) overnight. Cells were washed, re-suspended in normalization beads (Fluidigm), and acquired on a CyTOF2 cytometer (Fluidigm). Mass cytometry (CyTOF) data were analyzed using Cytobank. We used non-cardiomyocytes from individual hearts for each experimental replicate.

### Flow Cytometry

Isolated cardiac non-cardiomyocytes or cultured cardiac fibroblasts were stained with Zombie Aqua (1:200, Biolegend 423101) to discriminate between viable and dead cells. Cells were washed with cell staining buffer (Biolegend 420201) before blocking for non-specific binding with TruStain FcX (Biolegend 101320). Staining for cell surface markers was performed with fluorophore-conjugated primary antibodies (Major Resources Table) for 30 mins at 4C. Flow cytometry experiments included unstained cells and single-stained samples to visualize cells in the forward-side scatters and as compensation controls. All panels were validated using fluorescence minus one and isotype controls. For intracellular staining, BD Cytofix/Cytoperm kit (BD Bioscience 554714) was used to fix and permeabilize cells before staining against intracellular antigens, using fluorophore-conjugated antibodies (Major Resources Table). Flow cytometry data was acquired on a BD Fortessa X-20 or a Fortessa X-30 H0081 (BD Biosciences) cytometer and analyzed using FlowJo software.

### Fibroblast culture

Isolated cardiac non-cardiomyocytes were plated on non-coated dishes for 2 hours in DMEM supplemented with 10% FBS and 1% Penicillin/Streptomycin, followed by washing off non-adherent cells. Fibroblasts (adherent cells) were cultured until confluency, passaged, counted, and plated in 4- or 8-well ibiTreat micro slides (Ibidi 80426, 80806). The next day, fibroblasts were washed twice with PBS and cultured in DMEM supplemented with 1% Penicillin/Streptomycin without FBS. 5-8 hours later fibroblasts were stimulated with 100 ng/ml Recombinant CCL24 or (0.5 ug/mL) CCR3 antibody (MAB155) or 100 pg/ml of TGFb for 24-48h as indicated. For imaging, cells were fixed in 4% PFA for 15 minutes at room temperature, permeabilized and blocked in 0.3% Triton X-100 in PBS with 5% normal donkey serum and 1% BSA and stained for alpha Smooth Muscle Actin (aSMA, 1:500), Collagen-1 (1:240), periostin (1:200) and DAPI (1:2,000) in permeabilization buffer without donkey serum.

### Quantitative PCR

Heart tissue was harvested from mice and 20-30mg chunks were subjected to total RNA extraction using the RNeasy Plus Mini kit (Qiagen), and subsequent cDNA preparation was carried out with the iScript cDNA Synthesis kit (Bio-Rad). cDNA was quantified using SYBR Green Supermix (Bio-Rad) on a CFX Connect qPCR machine (Bio-Rad). Normalization of gene expression to GAPDH was performed, and changes in gene expression were determined using the 2(-ΔΔCT) method.

### Bone marrow chimera

Bone marrow cells were flushed out from the femurs and tibias of donor mice (WT or CCL24 KO). Recipient mice (WT) received a total lethal dose of 600 rads. Irradiations were done in 2 doses (300 rads each) separated by a period of 3-4 hours. Two hours after the second irradiation, the recipient mouse received at least 10×10^6^ of bone marrow donor cells through intravenous injection via tail vein. Bone marrow chimeras were used at 8 weeks after the introduction of bone marrow donor cells.

### Histology

Freshly harvested hearts were immediately fixed in 10% formalin and underwent histological assessment for cross-section myocyte size through wheat germ agglutinin (WGA) staining or fibrosis through Movat’s pentachrome and Picrosirius Red Fast Green staining by Biorepository & Laboratory Services. The fibrotic scar area fraction in images at 20X resolution was quantified using the iLastik Segmentation software and macros scripted in FIJI software (ImageJ). Fibrotic areas (red or yellow) were segmented from muscle and background using iLastik. The generated segmentation heatmaps were input to ImageJ, individual channels were split and scar area fraction was measured as total fibrotic area divided by total muscle area. The cardiomyocyte cross-sectional area was quantified from images taken with 20x Objective, where each heart was represented by 3 random images from the left ventricle. At least 50 cardiomyocytes were traced from each image using ImageJ.

### Statistical Analyses

Results are expressed as mean +/− SEM. Normality tests were conducted for all analyses using the Shapiro-Wilk test. For comparing two groups with normal distribution, p-values were determined using an unpaired, two-tailed t-test, while Mann-Whitney U tests were used for non-normal distribution. For experiments with three or more groups, one-way ANOVA with Fisher’s LSD test was performed to determine differences. Normal distribution (Shapiro-Wilk test) and equal variances (Brown-Forsythe) were confirmed for all datasets analyzed. Statistical analyses were conducted using GraphPad Prism software.

## REFERENCES

1. Ridker, P.M. et al. Low-Dose Methotrexate for the Prevention of Atherosclerotic Events. N Engl J Med 380, 752–762 (2019).

2. Ridker, P.M. et al. Antiinflammatory Therapy with Canakinumab for Atherosclerotic Disease. N Engl J Med 377, 1119–1131 (2017).

3. Hulsmans, M. et al. Macrophages Facilitate Electrical Conduction in the Heart. Cell 169, 510–522.e520 (2017).

4. Nicolás-Ávila, J.A. et al. A Network of Macrophages Supports Mitochondrial Homeostasis in the Heart. Cell 183, 94–109.e123 (2020).

5. Epelman, S. et al. Embryonic and adult-derived resident cardiac macrophages are maintained through distinct mechanisms at steady state and during inflammation. Immunity 40, 91–104 (2014).

6. Bajpai, G. et al. Tissue Resident CCR2− and CCR2+ Cardiac Macrophages Differentially Orchestrate Monocyte Recruitment and Fate Specification Following Myocardial Injury. Circulation Research 124, 263–278 (2019).

7. Wong, N.R. et al. Resident cardiac macrophages mediate adaptive myocardial remodeling. Immunity 54, 2072–2088.e2077 (2021).

8. Zaman, R. et al. Selective loss of resident macrophage-derived insulin-like growth factor-1 abolishes adaptive cardiac growth to stress. Immunity 54, 2057–2071.e2056 (2021).

9. Hulsmans, M. et al. Cardiac macrophages promote diastolic dysfunction. J Exp Med 215, 423–440 (2018).

10. Forssmann, U. et al. Eotaxin-2, a Novel CC Chemokine that Is Selective for the Chemokine Receptor CCR3, and Acts Like Eotaxin on Human Eosinophil and Basophil Leukocytes. Journal of Experimental Medicine 185, 2171–2176 (1997).

11. Greenman, R., et al. CCL24 regulates biliary inflammation and fibrosis in primary sclerosing cholangitis. JCI Insight 8 (2023).

12. Segal-Salto, M. et al. A blocking monoclonal antibody to CCL24 alleviates liver fibrosis and inflammation in experimental models of liver damage. JHEP Rep 2, 100064 (2020).

13. Diny, N.L. et al. Macrophages and cardiac fibroblasts are the main producers of eotaxins and regulate eosinophil trafficking to the heart. Eur J Immunol 46, 2749–2760 (2016).

14. Wang, Z. et al. Mechanistic basis of neonatal heart regeneration revealed by transcriptome and histone modification profiling. Proc Natl Acad Sci U S A 116, 18455–18465 (2019).

15. Wang, Z. et al. CCL24/CCR3 axis plays a central role in angiotensin II–induced heart failure by stimulating M2 macrophage polarization and fibroblast activation. Cell Biology and Toxicology 39, 1413–1431 (2023).

16. Revelo, X.S. et al. Cardiac Resident Macrophages Prevent Fibrosis and Stimulate Angiogenesis. Circ Res 129, 1086–1101 (2021).

17. Lee, S.H. et al. M2-like, dermal macrophages are maintained via IL-4/CCL24–mediated cooperative interaction with eosinophils in cutaneous leishmaniasis. Science Immunology 5, eaaz4415 (2020).

18. Makita, N., Hizukuri, Y., Yamashiro, K., Murakawa, M. & Hayashi, Y. IL-10 enhances the phenotype of M2 macrophages induced by IL-4 and confers the ability to increase eosinophil migration. Int Immunol 27, 131–141 (2015).

19. Froese, N. et al. Analysis of myocardial cellular gene expression during pressure overload reveals matrix based functional intercellular communication. iScience 25, 103965 (2022).

20. Filippone, R.T. et al. Potent CCR3 Receptor Antagonist, SB328437, Suppresses Colonic Eosinophil Chemotaxis and Inflammation in the Winnie Murine Model of Spontaneous Chronic Colitis. Int J Mol Sci 23 (2022).

21. Aguado-Alvaro, L.P. et al. Comparative Evaluation of Inducible Cre Mouse Models for Fibroblast Targeting in the Healthy and Infarcted Myocardium. Biomedicines 10 (2022).

22. Saul, D. et al. A new gene set identifies senescent cells and predicts senescence-associated pathways across tissues. Nature Communications 13, 4827 (2022).

23. Katakura, Y., Totsuka, M., Imabayashi, E., Matsuda, H. & Hisatsune, T. Anserine/Carnosine Supplementation Suppresses the Expression of the Inflammatory Chemokine CCL24 in Peripheral Blood Mononuclear Cells from Elderly People. Nutrients 9 (2017).

24. Ma, Y. et al. Deriving a cardiac ageing signature to reveal MMP-9-dependent inflammatory signalling in senescence. Cardiovasc Res 106, 421–431 (2015).

25. Kanellakis, P., Ditiatkovski, M., Kostolias, G. & Bobik, A. A pro-fibrotic role for interleukin-4 in cardiac pressure overload. Cardiovascular Research 95, 77–85 (2012).

26. Peng, H. et al. Profibrotic Role for Interleukin-4 in Cardiac Remodeling and Dysfunction. Hypertension 66, 582–589 (2015).

27. Gieseck, R.L., Wilson, M.S. & Wynn, T.A. Type 2 immunity in tissue repair and fibrosis. Nature Reviews Immunology 18, 62–76 (2018).

28. Van Dyken, S.J. & Locksley, R.M. Interleukin-4- and interleukin-13-mediated alternatively activated macrophages: roles in homeostasis and disease. Annu Rev Immunol 31, 317–343 (2013).

29. Travers, J.G., Kamal, F.A., Robbins, J., Yutzey, K.E. & Blaxall, B.C. Cardiac Fibrosis. Circulation Research 118, 1021–1040 (2016).

30. Haider, N. et al. Transition of Macrophages to Fibroblast-Like Cells in Healing Myocardial Infarction. J Am Coll Cardiol 74, 3124–3135 (2019).

31. Frangogiannis, N.G. Cardiac fibrosis. Cardiovasc Res 117, 1450–1488 (2021).

32. Hu, S. et al. Different Roles of Resident and Non-resident Macrophages in Cardiac Fibrosis. Front Cardiovasc Med 9, 818188 (2022).

33. Kain, D. et al. Macrophages dictate the progression and manifestation of hypertensive heart disease. International Journal of Cardiology 203, 381–395 (2016).

34. Savill, J. Apoptosis in resolution of inflammation. Journal of Leukocyte Biology 61, 375–380 (1997).

35. Huynh, M.-L.N., Fadok, V.A. & Henson, P.M. Phosphatidylserine-dependent ingestion of apoptotic cells promotes TGF-β1 secretion and the resolution of inflammation. The Journal of Clinical Investigation 109, 41–50 (2002).

36. Matthews, A.N. et al. Eotaxin is required for the baseline level of tissue eosinophils. Proc Natl Acad Sci U S A 95, 6273–6278 (1998).

37. Kitaura, M. et al. Molecular cloning of human eotaxin, an eosinophil-selective CC chemokine, and identification of a specific eosinophil eotaxin receptor, CC chemokine receptor 3. J Biol Chem 271, 7725–7730 (1996).

38. Tannu, S., Allocco, J., Yarde, M., Wong, P. & Ma, X. Experimental model of congestive heart failure induced by transverse aortic constriction in BALB/c mice. Journal of Pharmacological and Toxicological Methods 106, 106935 (2020).

39. Subramanian, A. et al. Gene set enrichment analysis: a knowledge-based approach for interpreting genome-wide expression profiles. Proc Natl Acad Sci U S A 102, 15545–15550 (2005).

