## Supplemental Figure 1 for "Macrophage-derived CCL24 promotes fibrosis and worsens cardiac dysfunction during heart failure"

**Supplementary Figure 1**

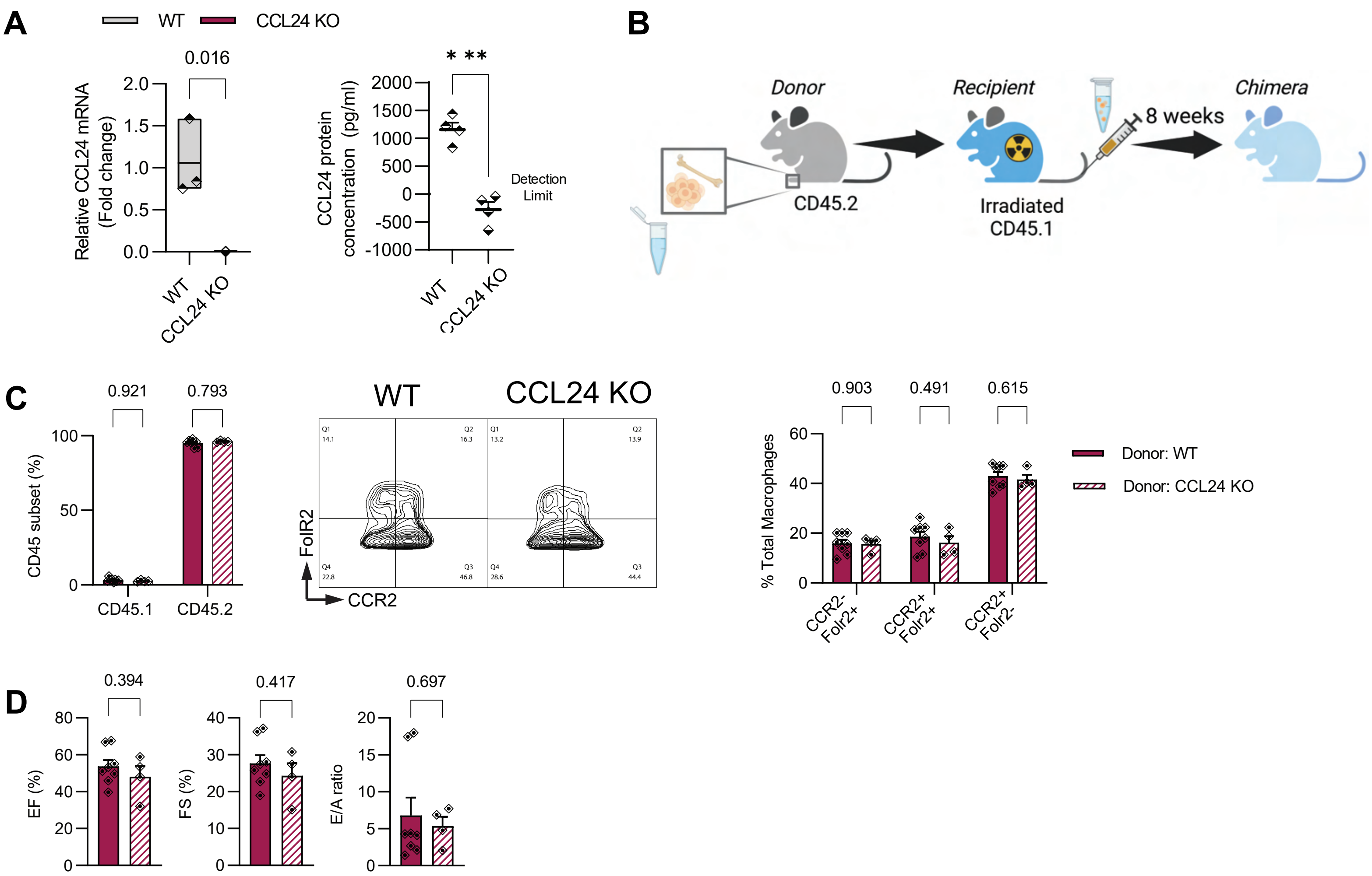

**Supplementary Figure 2**

A

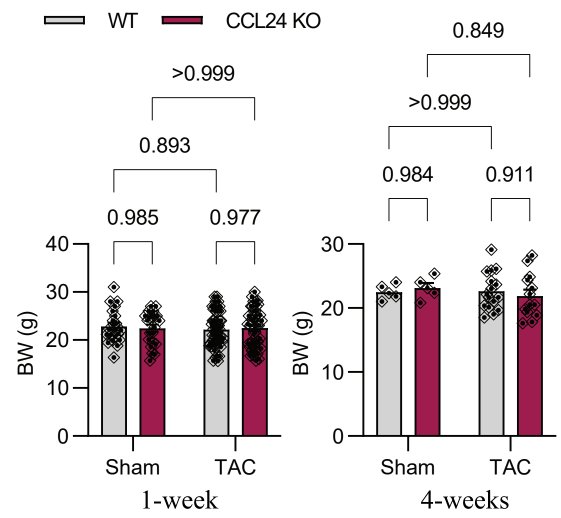


B

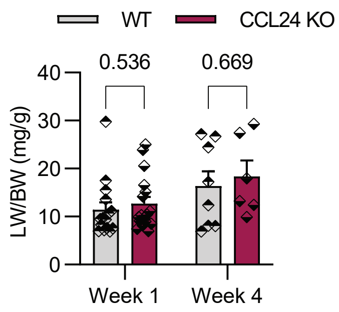


C

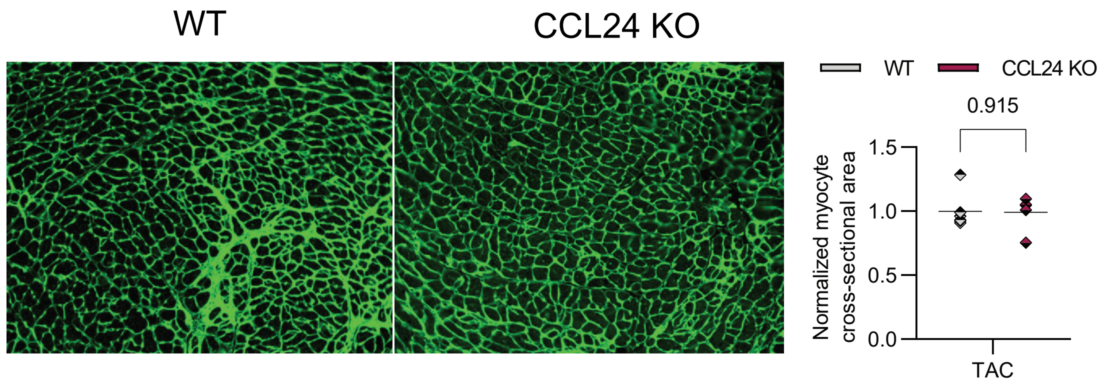

**Supplementary Figure 3**

**A**

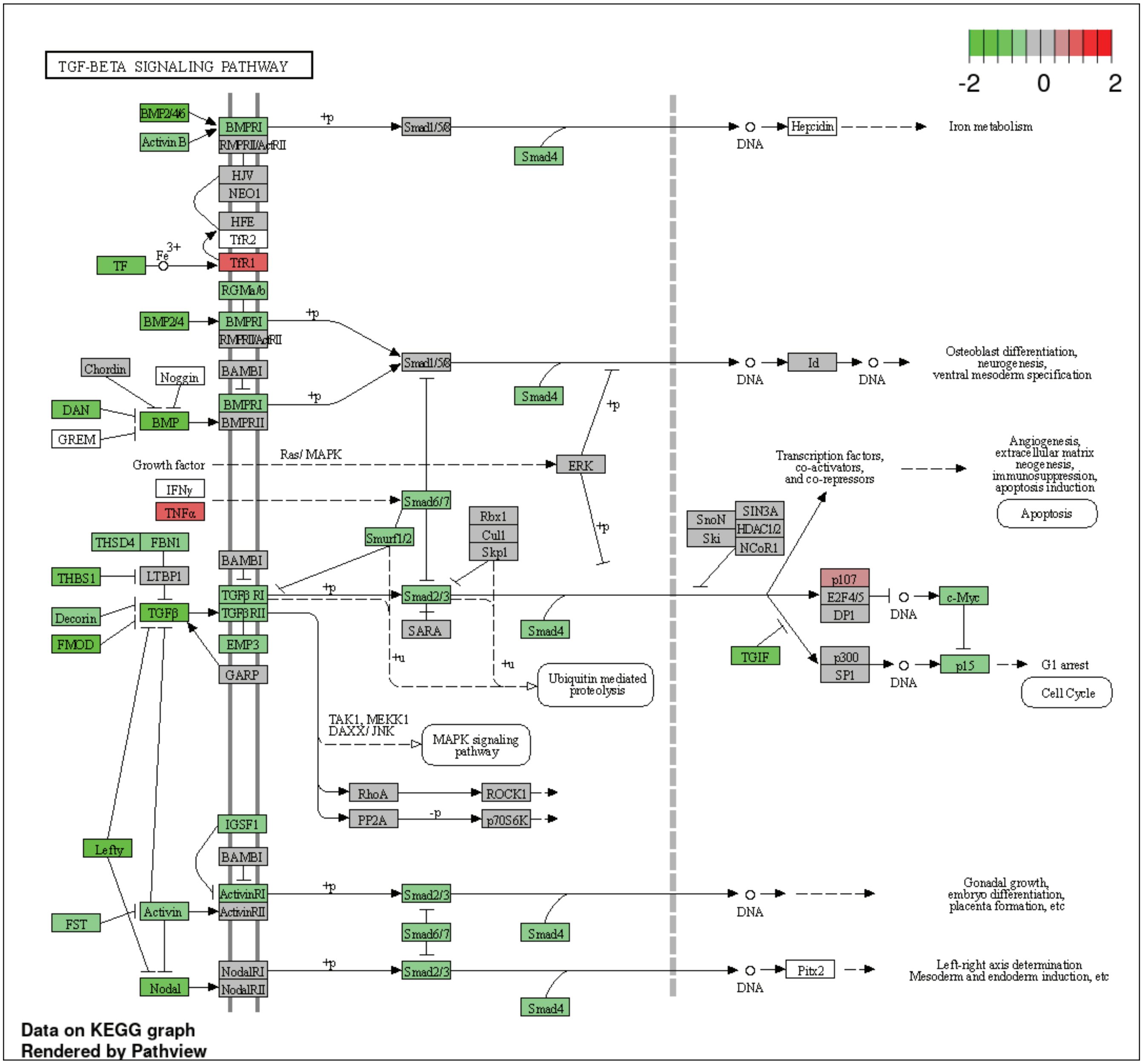

**Supplementary Figure 4**

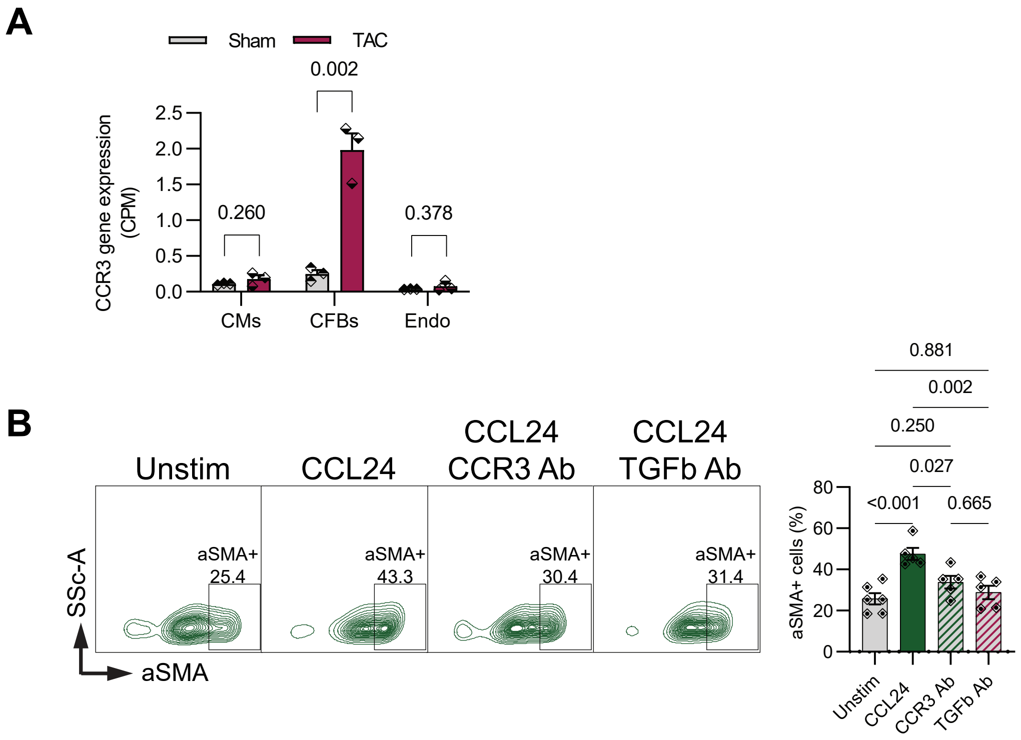

**Supplementary Figure 5**

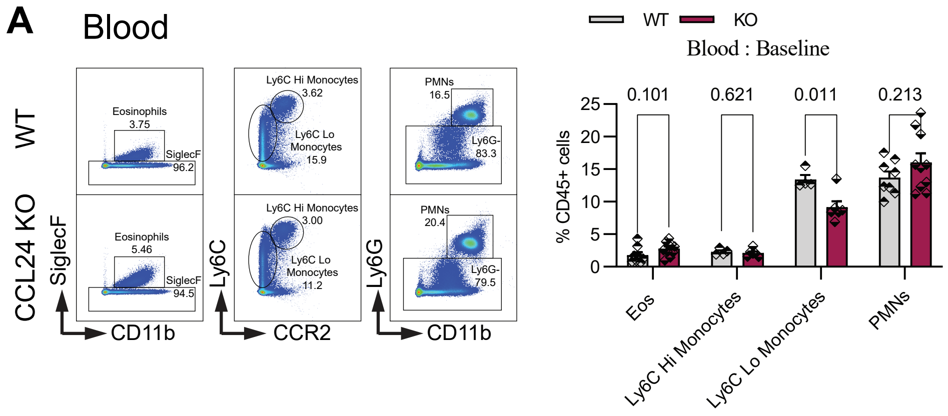


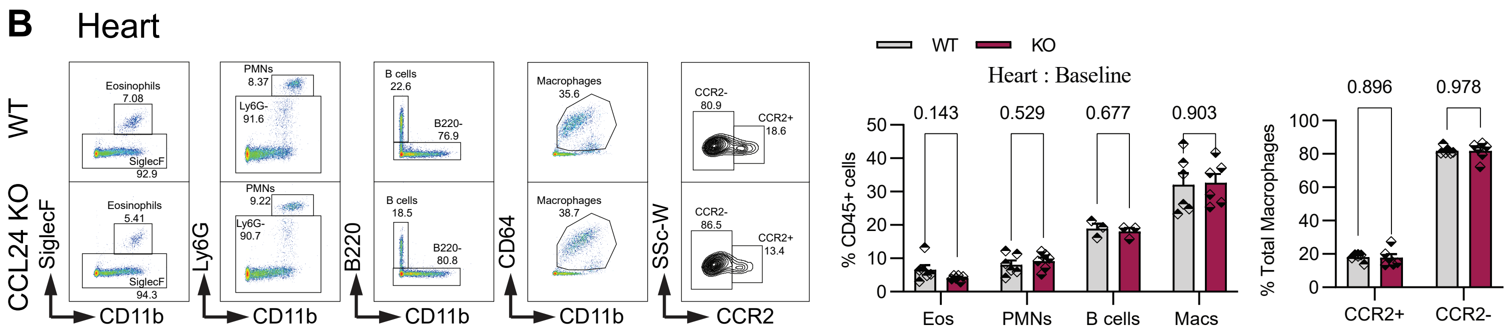
